# High-density linkage map and QTLs for growth in snapper (*Chrysophrys auratus*)

**DOI:** 10.1101/376012

**Authors:** David T. Ashton, Peter A. Ritchie, Maren Wellenreuther

**Affiliations:** The New Zealand Institute for Plant & Food Research Limited, Nelson, New Zealand; School of Biological Sciences, Victoria University of Wellington, Wellington, New Zealand

**Keywords:** linkage map, genome, fish, QTLs, growth, genotyping-by-sequencing

## Abstract

Characterizing the genetic variation underlying phenotypic traits is a central objective in biological research. This research has been hampered in the past by the limited genomic resources available for most non-model species. However, recent advances in sequencing technology and related genotyping methods are rapidly changing this. Here we report the use of genome-wide SNP data from the ecologically and commercially important marine fish species *Chrysophrys auratus* (snapper) to 1) construct the first linkage map for this species, 2) scan for growth QTLs, and 3) search for candidate genes in the surrounding QTL regions. The newly constructed linkage map contained ~11K SNP markers and is the densest map to date in the fish family Sparidae. Comparisons with available genome scaffolds indicated that overall marker placement was strongly correlated between the scaffolds and linkage map (R = 0.7), but at fine scales (< 5 cM) there were some precision limitations. Of the 24 linkage groups, which reflect the 24 chromosomes of this species, three were found to contain QTLs with genome-wide significance for growth-related traits. A scan for 13 known candidate growth genes located the genes for growth hormone, parvalbumin, and myogenin within 13.2, 2.6, and 5.0 cM of these genome-wide significant QTLs, respectively. The linkage map and QTLs found in this study will advance the investigation of genome structure and selective breeding in snapper.

## INTRODUCTION

Characterizing the genetic variation that affects phenotypic traits is a central goal in biology. Understanding this variation can inform selective breeding programmes (Dekkers, 2012), be used to predict disease risk in medicine (Lehner, 2013), and help researchers to understand evolution in natural populations (Savolainen et al., 2013). While genetic research has typically been pioneered in laboratory model species, the development of cheaper high-throughput genomic methods (e.g. next generation sequencing) is now allowing this research to be extended to a wide range of non-model species (Braasch et al., 2015; Hilario, 2015).

Locating and characterizing quantitative trait loci (QTLs) is one common approach used to investigate how genetic variation influences a specific phenotype (e.g. Barria et al., 2017; Chen et al., 2017; Pértille et al., 2017). QTL mapping methods locate specific loci that are influencing a phenotypic trait based on a significant correlation between allelic variation of that loci and variation of the trait (Lynch and Walsh, 1998). New QTLs can be informative as standalone observations, or used to identify candidate genes and causative variants in the surrounding genome, which may be influencing the trait (e.g. Bettembourg et al., 2017). Genotype-phenotype datasets can also be used to develop multi-marker models (based on multiple QTLs) which explain variation of one or more phenotypic traits (e.g. genomic selection models) (Cros et al., 2017).

Having a road-map of the genome (e.g. linkage map or high quality genome assembly) is an important prerequisite for QTL mapping, as it allows the relative positioning of different loci. High-quality genome assemblies are most effective because they allow genetic markers to be positioned at a base-pair level while also providing sequence information for the surrounding area. However, most non-model species do not currently have chromosome-level genome assemblies and instead rely on linkage maps to ascertain the relative position of markers in the genome (Braasch et al., 2015). Although they are much less precise than high quality genome assemblies, linkage maps can serve the dual purpose of bridging the resource gap before a genome is developed and providing useful information to improve the arrangement of scaffolds during the genome assembly process (Fierst, 2015). Data sets that are developed for constructing a linkage map can also be used in QTL mapping.

Research on marine teleost fish is one area that has potential to benefit from more affordable and higher-throughput genomic technologies. Teleost fish are the largest group of vertebrates, with over 26,000 species of fish (Miller and Harley, 2006), most which have limited geographic range and many are of commercial significance. The high diversity of production species means that scientific efforts are spread thinly, with few to no genomic resources available for many species. One such species, and the focus of this study, is the marine finfish *Chrysophrys auratus* (henceforth referred to as “snapper’). Snapper supports a valuable recreational and commercial inshore fishery around the northern parts of New Zealand, southern Australia, and some of the Pacific Islands and is a strong candidate for development into an aquaculture species in both New Zealand and Australia. However, until recently there has been limited or no genomic research carried out on this species One of the most important traits in farmed species is growth rate, as it directly affects the efficiency of production systems. In many animal species growth is a complex trait that is influenced by a network of genes (De-Santis and Jerry, 2007) and many environmental factors, such as seasonal variation in temperature, food availability, and competition (Handeland et al., 2008). Moreover, growth in fish is also known to be correlated with variation in other life-history traits, such as gonad maturation processes and reproductive timing (Bhatta et al., 2012; Park et al., 2016). Despite the numerous factors influencing growth, most studies that investigate growth report moderate to high heritability (e.g. 0.1 - 0.5) throughout a wide range of taxa (Tsai et al., 2015; Wang, 2009; Ye et al., 2017). A number of genes associated with growth rate in fish have been identified by other means (reviewed in De-Santis and Jerry, 2007) including growth hormone, insulin-like growth factors, and a range of myogenic growth regulators.

This study follows on from previous work that investigated the pedigree structure, inbreeding, and heritability of traits in a new population of snapper (Ashton et al., 2018). The primary goal was to identify genetic variation underlying growth rate, which can be used to inform a newly started selective breeding programme. Here we use genome-wide SNP data generated in this original study to 1) construct a high density linkage map, 2) conduct QTL mapping for three measures of growth rate (peduncle length, fork length, and weight), and 3) investigate the position of 13 candidate growth genes and their relative position to growth rate QTLs.

## MATERIALS AND METHODS

### Study population

A snapper breeding programme is under development at The New Zealand Institute for Plant & Food Research Limited and currently includes a population with three generations held at the Nelson Research Centre (Ashton et al., 2018). Data from the two most recent generations (F_1_ = 70 individuals, F_2_ = 577 individuals) were used in the current study. Uncontrolled tank-based spawning of the F_1_ generation in a single tank was used to produce the offspring F_2_ generation. This resulted in a complex pedigree, meaning that we obtained a combination of full-sibling, half-sibling, and unrelated individuals in the F_2_ generation (Ashton et al., 2018). The F_2_ offspring were held in a single tank until they were approximately one year old and then split evenly among four tanks with comparable feeding, light, water flow, aeration, tank design. All research carried out in this study was reviewed and approved by the animal ethics committee of Victoria University of Wellington (Application number 2014R19).

### Phenotyping

The three measures of growth rate used in the current study were fork length, peduncle length, and weight. Fork length was the distance from the nose to the fork in the tail. Peduncle length was the distance from the nose to the narrowest cross-section across the tail. Measurements were made when the fish were approximately one year old (436-487 days) and again when they were approximately three years old (1045-1131 days). Length measurements were made by collecting images of each individual and then making measurements from those images. A ruler was included in each image to provide a scale. The number of individuals measured differed between year one and year three as a result of natural mortality during the study.

### Preparation of GBS libraries

Samples of fin tissue were collected for all fish and DNA was extracted from these samples using a modified salt extraction protocol (Ashton et al., 2018). DNA was genotyped using a modified genotyping by sequencing approach (Hilario, 2015). A total of eight pooled libraries were sequenced on an Illumina HiSeq 2500. Each pool contained DNA from 96 individuals which were individually barcoded. Duplicate or triplicate samples were prepared for each of the parents and grandparents, and single samples for each of the offspring. Sequencing data from the GBS libraries were processed using the STACKs pipeline (Catchen et al., 2013) and the genotyping results exported to a GENEPOP format txt file. When variant calling, sequence stacks were aligned to an initial snapper genome sequence (Wellenreuther et al., unpublished) in the STACKs pipeline. SNPs from the STACKs pipeline were filtered for > 7x coverage, presence in >75% of the population, and a minor allele frequency > 0.05.

### Linkage map construction

The parents for each individual in the dataset were previously identified using Cervus (Kalinowski et al., 2007) in Ashton et al. (2018). A linkage map was constructed based on the genotyping and pedigree data in LEPMAP 2.0 (Rastas et al., 2016). Data from the largest 14 F_2_ families (full and half-sibling families) were used, and included a total of 269 offspring and 14 parents. A total of 20,311 SNPs were located in the initial genotyping file. Markers were separated into chromosomes with the SeparateChromosomes module (logarithm of odds (LOD) limit = 14, minimum markers per linkage group = 50). The best order was then generated with the OrderMarkers module. Markers near the start and end of each linkage group (start and end 10% based on centimorgan (cM) distance) were removed if they were more than 3 cM from the next closest marker. The accuracy and precision of the final linkage map was reviewed by comparing the linkage map position (cM) with the position of markers on available genome scaffolds (base-pairs) from the snapper genome. This was possible by using the STACKs output files, which showed the scaffolds on which each marker was placed as well as their bp position. The correlation between marker position in cM and bp position on the scaffolds was calculated for all scaffolds that contained >50 SNPs. Whether scaffolds were placed uniquely on one of the 24 expected linkage groups was also investigated. The extent of linkage and linkage disequilibrium across the linkage groups was reviewed by calculating the pairwise linkage disequilibrium results for each set of markers in PLINK 1.9 (Purcell et al., 2007) and then visualizing the mean value at different distances across the linkage groups in R statistical environment (version: 3.2.3) (R Core Team, 2013). This was done for all individuals in the F_2_ generation and separately for individuals in the largest full sibling F_2_ family. The sex-specific recombination rate was calculated by comparing the linkage map distance (cM) and genome scaffold distance (bp) between individual marker pairs for males and females.

### QTL identification

Quantitative trait loci identification was carried out using QTDT 2.6.1 (Abecasis et al., 2000). This software performs a transmission disequilibrium test based on the differential transfer of alleles from parents to offspring (i.e. parents act as controls for population stratification). With parents as controls, transmission disequilibrium tests are unaffected by complex pedigree sample designs (Spielman and Ewens, 1996). Genotype data from the F_1_ and F_2_ generations and phenotyping data from the F_2_ generation were used. Only markers that had been placed on the linkage map were included in this analysis. Before running the analysis, the genotype data were filtered for Mendelian errors by dropping loci for any individual that contained alleles not observed in either of the two parents. The phenotype measurements used for the analysis were standardized by tank and date collected to correct for temporal and tank effects. The QTL scan results from QTDT were visualised using the ggplot2 library in the R statistical environment (version: 3.2.3) (R Core Team, 2013). A Bonferroni correction was used to calculate 95% confidence limits (i.e. 0.05 / number of markers) at a linkage group and genome-wide scale. A randomized dataset was also put through QTDT 2.6.1 to check for the possibility of false positive QTL signals.

This dataset was constructed by randomizing the phenotypes between offspring within each family, but leaving the genotype data untouched.

### Candidate genes and their location

The position of 13 candidate growth genes for fish (De-Santis and Jerry, 2007) and QTLs for growth identified in this study were compared using an initial snapper reference genome (Wellenreuther et al., unpublished). To do this, the sequence for each candidate gene was located on the NCBIS nucleotide database from the closest related teleost species - either the DNA or mRNA sequence. DNA sequences were aligned with the genome scaffolds by selecting the largest exon for the target gene and aligning with the “Map to Reference” alignment using the “Geneious mapper” in Geneious 10.0.9 (Kearse et al., 2012); alignment sensitivity was set to “High Sensitivity / Medium” with default settings. For mRNA sequences the sequences were aligned with the “Map to Reference” alignment using the “RNA Seq” mapper in Geneious 10.0.9; alignment sensitivity was set to “High Sensitivity / Medium” with the maximum gap size increased to 1000 bp. For each alignment the percentage of matching base pairs was reported for the largest exon. The linkage group and cM position of the scaffold containing specific candidate genes was then located using the STACKs output files and the newly constructed linkage map. Sequence data are available at GenBank; trait and linkage map data at Zenodo, the accession numbers are listed in Supplementary table 3 (this will be added upon acceptance of this paper).

## RESULTS

### Phenotyping

Peduncle length, fork length, and weight were recorded when individuals where 436-487 days old (Year one) and 1045-1131 days old (Year three). The distribution and relative sizes of fish in year one and year three are illustrated in Figure 1. In the first set of measurements the mean and standard deviation for fork length, peduncle length, and weight were 160.1 ± 15.0 mm, 132.1 ± 12.3 mm, and 89.8 ± 24.3 g, respectively (Table 1). In the second set of measurements the same measures were 257.7 ± 21.0, 214.3 ± 17.0, and 363.2 ± 84.9, respectively (Table 1). The three measures for growth were all found to be strongly positively correlated (Pearson’s R > 0.93, Ashton et al., 2018). Strong positive correlation was also observed between year one and year three for each measure (Pearson’s R = 0.71 – 0.73; Supplementary figure 1).

**Table 1.** Peduncle length, fork length, and weight at year one and year three including number or measurements (n), mean, and standard deviation (stdev).

|  | Year one |  |  | Year three |  |  |
| --- | --- | --- | --- | --- | --- | --- |
|  | n | mean | stdev | n | mean | stdev |
| **Peduncle length (mm)** | 768 | 132.1 | 12.3 | 417 | 214.3 | 17 |
| **Fork length (mm)** | 768 | 160.1 | 15.0 | 417 | 257.7 | 21.0 |
| **Weight (g)** | 762 | 89.8 | 24.3 | 324 | 363.2 | 84.9 |

**Figure 1.**
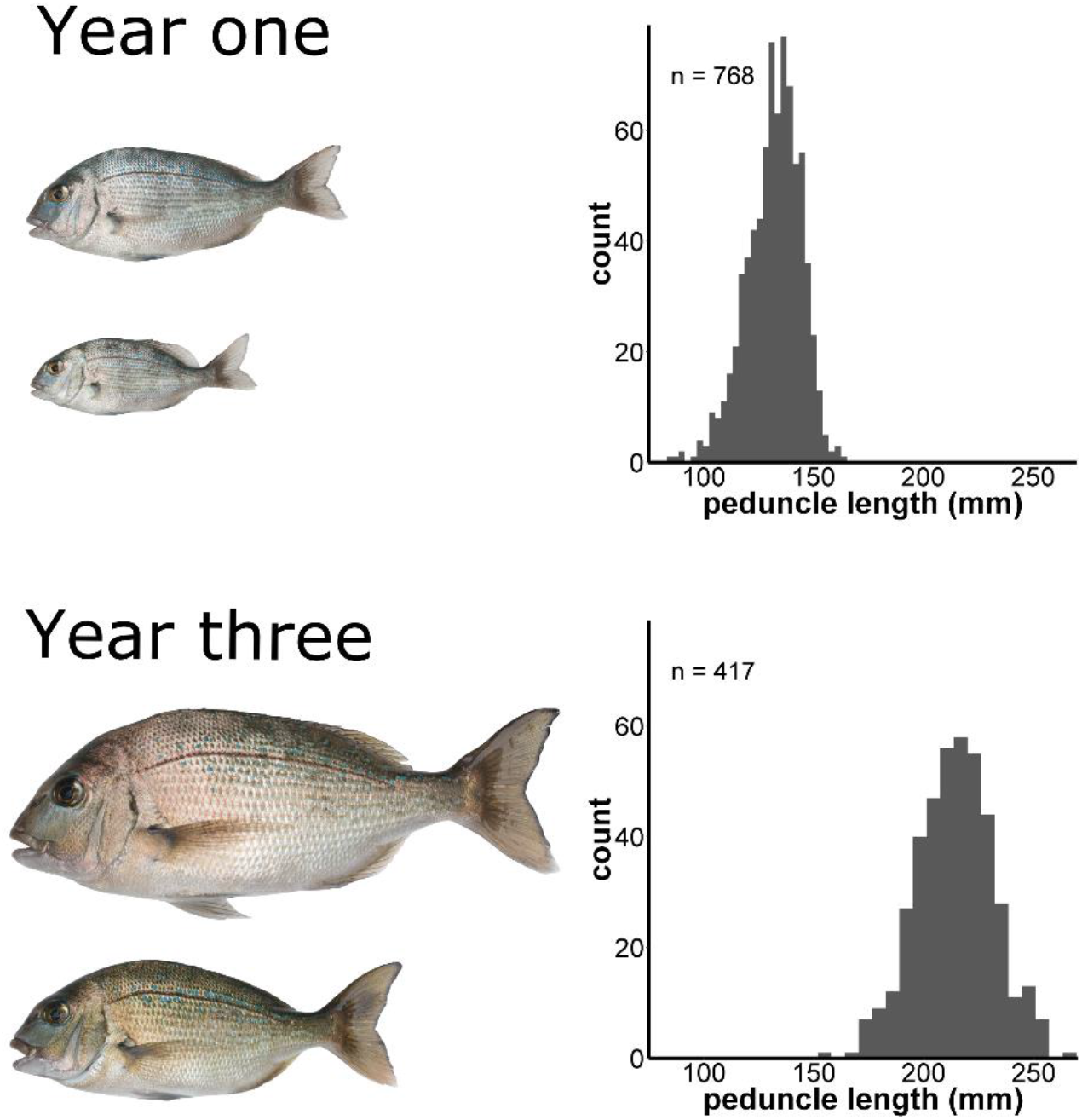
Peduncle length measurements at year one and year three. Images on the left side of the diagram are to scale relative to each other and show the smallest and largest individual fish from a tank of F_2_ individuals at year one and year three, respectively.

### Genotyping by sequencing

A total of 1.6 billion reads were produced for all eight pooled GBS libraries with approximately 2, 4 or 6 million reads for each single, duplicate, or triplicate individual library respectively. Using the STACKs pipeline a total of 20,311 SNPs were found after filtering for >7x coverage, present in 75% of the individuals in the population, and a minor allele frequency (MAF) of 0.05.

### Pedigree structure and linkage map

Parents were identified for 93% of the individuals in the F_1_ and F_2_ generations. The remaining 7% with missing parents were mainly located in the F_1_ generation, and were the result missing F_0_ wild-caught individuals that were not available for sampling at the time of the study. A mixture of full sibling and half sibling families was present in the F_2_ generation.

A total of 10,968 SNPs were positioned on the linkage map (Figure 2). The total length of the sex-averaged linkage map was 1,363.0 cM with an average marker spacing of 0.129 cM. The lengths of the male and female maps were 1401.5 cM and 1359.0 cM, respectively. The female and male recombination rates were 3.28 cM/Mb and 1.93 cM/Mb based on comparison with available scaffolds from the initial snapper genome (Wellenreuther et al., unpublished). When aligning the genome scaffolds to the linkage map, individual scaffolds were placed exclusively onto one of the 24 linkage groups. Visualization of the linkage map and scaffold position for the four largest scaffolds showed a clear relationship between the ordering of markers (correlation = 0.91, 0.53, 0.78, and -0.95), but that some noise was apparent around the exact placement (Figure 3). The correlation for all scaffolds with 50 or more markers was similar (R = 0.74 ± 0.20 for 1723 markers on 26 scaffolds). The 95% confidence interval of the residuals ranged from -5.7 to 3.2 centimorgans with a mean of -1.25; indicating that markers were placed within ~4.5 cM of their correct location. Investigation of the degrees of linkage and linkage disequilibrium within the dataset showed a clear pattern of linkage decay over the length of the linkage groups (Figure 4). When looking at a single F_2_ family we see a high degree of linkage. However, when looking at the whole F_2_ generation the decay of linkage and linkage disequilibrium is much greater, with minimal linkage observed even over small distances.

**Figure 2.**
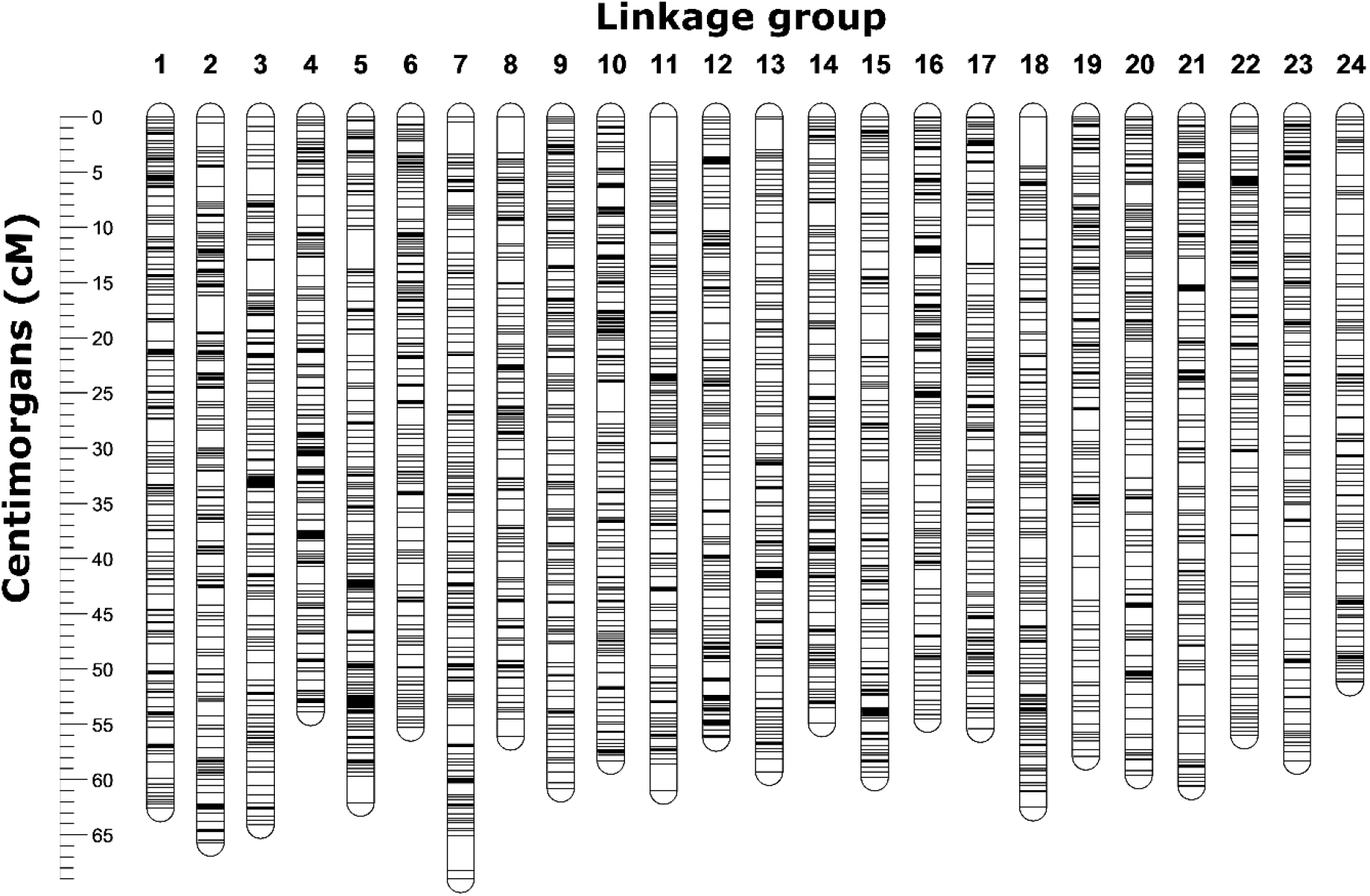
Visualization of the linkage map including a total of 10,968 SNPs placed along the map in 24 linkage groups. The 24 linkage groups are equivalent size and number to represent the expected 24 *Chyrsophrys auratus* chromosomes.

**Figure 3.**
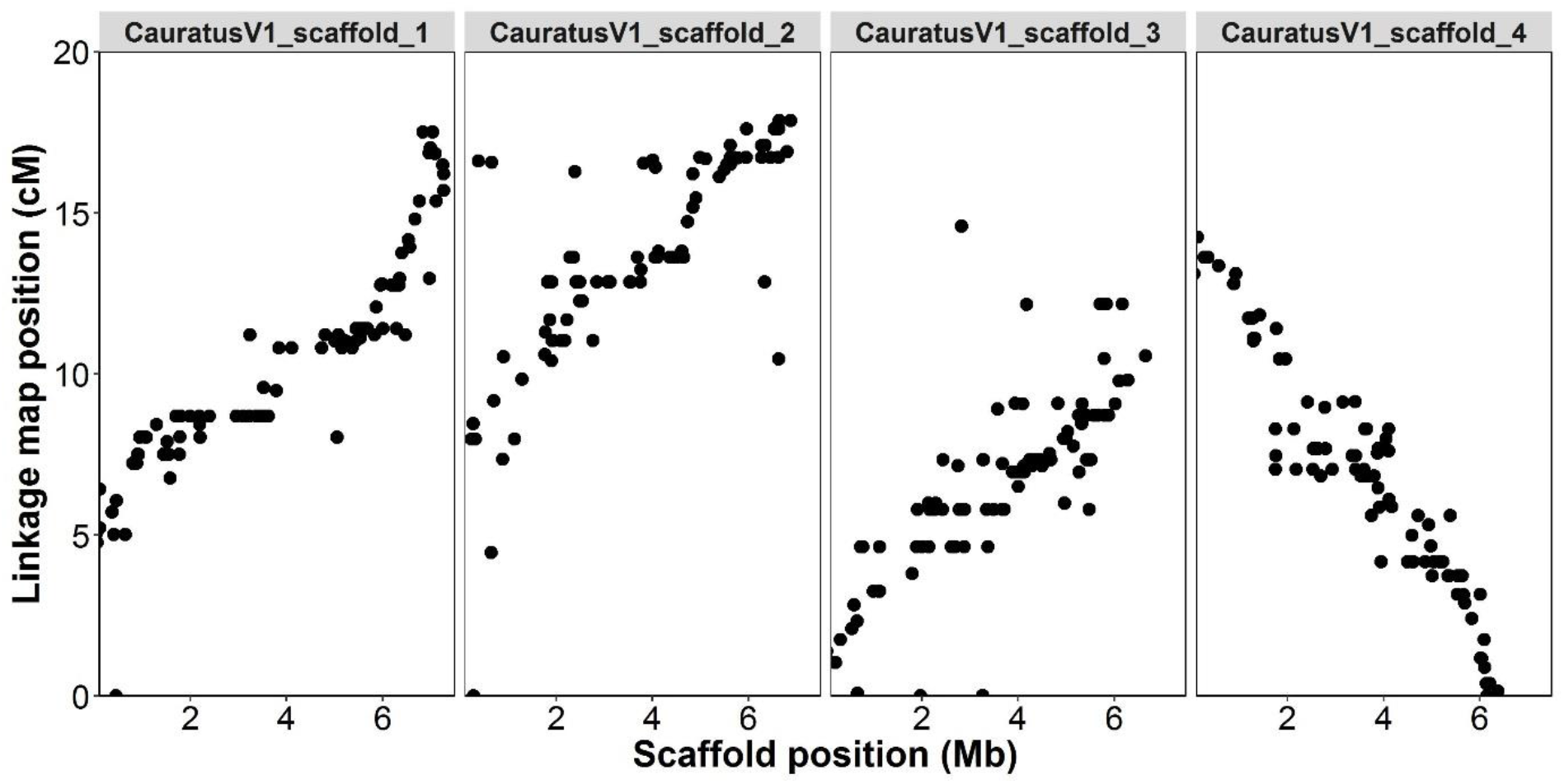
Comparison of linkage group (cM) and genome scaffold (Mb) position for loci placed on the four largest scaffolds available from the genome. The position was strongly correlated between the two approaches (scaffold_1 = 0.91, scaffold_2 = 0.53, scaffold_3 = 0.78, scaffold_4 = -0.95), but there is also noise around the precise placement on the linkage map.

**Figure 4.**
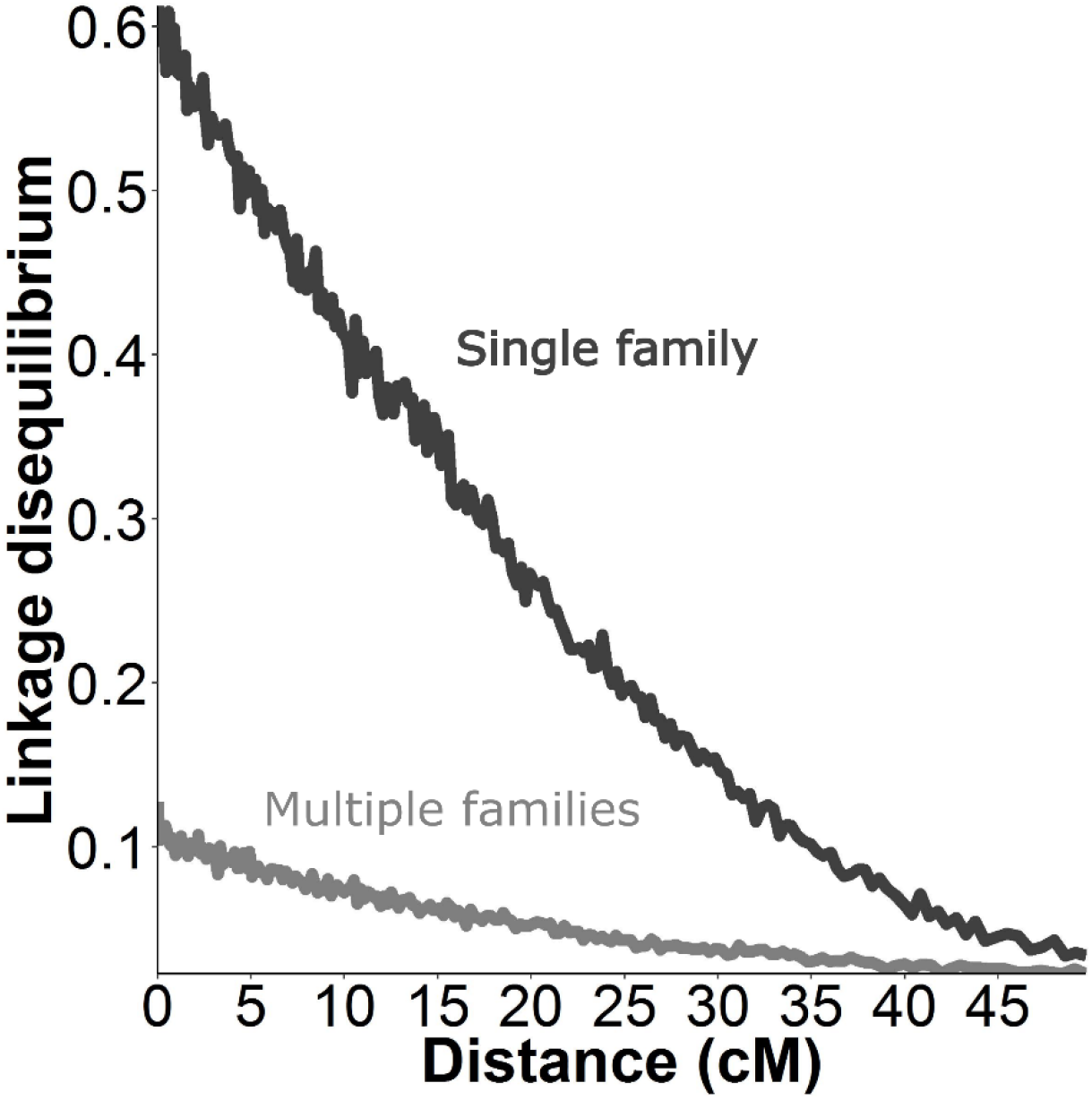
Linkage (R_2_ correlation) between each pair of markers in the dataset plotted against the distance (cM) between those markers on the linkage map. This statistic was calculated for the entire QTL mapping dataset (multiple families, families = 137, n = 539) and for the data from the single largest family in the dataset (n = 48). The results show a consistent decay of linkage across the length of the linkage maps and > 5x higher linkage in the largest family than the entire dataset.

### QTL mapping

Multiple QTLs were found for all three traits (Figure 5, Table 2, and Supplementary table 1). Most of the QTLs were shared among the three traits. QTLs on linkage groups 3, 11, and 16 passed the genome-wide significant –log10(p) of 5.33 (Figure 5 and Table 2). The QTLs on linkage group 3 and 11 were located in a small region (< 11 cM), but on linkage group 16 QTL peaks passing the genome-wide significance level were observed across the length of the linkage group. The QTLs found in year one had moderate effect sizes ranging from a minimum R^2^ of 0.04 to a maximum of 0.10. Genome-wide significant QTLs were not found for any of the traits in year three, but chromosome-wide significant QTLs were found on linkage groups 5, 11, 12, and 24. No false positive QTL signals were detected when the randomized dataset was put through the QTL mapping software.

**Table 2.** Putative QTLs that were significant for at least one trait at a genome-wide significance level of 5.33. Effect size (R^2^) was estimated in QTDT as the difference between the R squared values of the total model and the genotype model. Loci are reported including their linkage group number (LG) and position (cM). The linkage group-wide significance level is reported in the last column (LGW).

| LG | cM | loci | Fork length |  | Peduncle length |  | Weight |  | LGW |
| --- | --- | --- | --- | --- | --- | --- | --- | --- | --- |
|  |  |  | -log10(p) | R<sup>2</sup> | -log10(p) | R<sup>2</sup> | -log10(p) | R<sup>2</sup> | -log10(p) |
| 3 | 12.2 | 29323_42 | 6.40 | 0.09 | 6.10 | 0.08 | 4.52 | 0.06 | 3.98 |
| 11 | 1.7 | 17817_58 | 4.70 | 0.04 | 5.05 | 0.04 | 5.52 | 0.04 | 3.96 |
| 11 | 2.7 | 43276_58 | 4.52 | 0.04 | 4.40 | 0.04 | 5.22 | 0.05 | 3.96 |
| 11 | 10.0 | 99093_35 | 5.52 | 0.04 | 5.70 | 0.04 | 5.52 | 0.04 | 3.96 |
| 11 | 12.2 | 38917_81 | 5.30 | 0.07 | 5.70 | 0.08 | 5.52 | 0.08 | 3.96 |
| 16 | 11.0 | 80388_28 | 5.70 | 0.1 | 5.70 | 0.1 | 5.22 | 0.09 | 3.93 |
| 16 | 21.2 | 17603_14 | 5.52 | 0.07 | 5.15 | 0.06 | 4.52 | 0.05 | 3.93 |
| 16 | 40.2 | 39092_27 | 5.70 | 0.05 | 5.52 | 0.06 | 5.00 | 0.05 | 3.93 |

**Figure 5.**
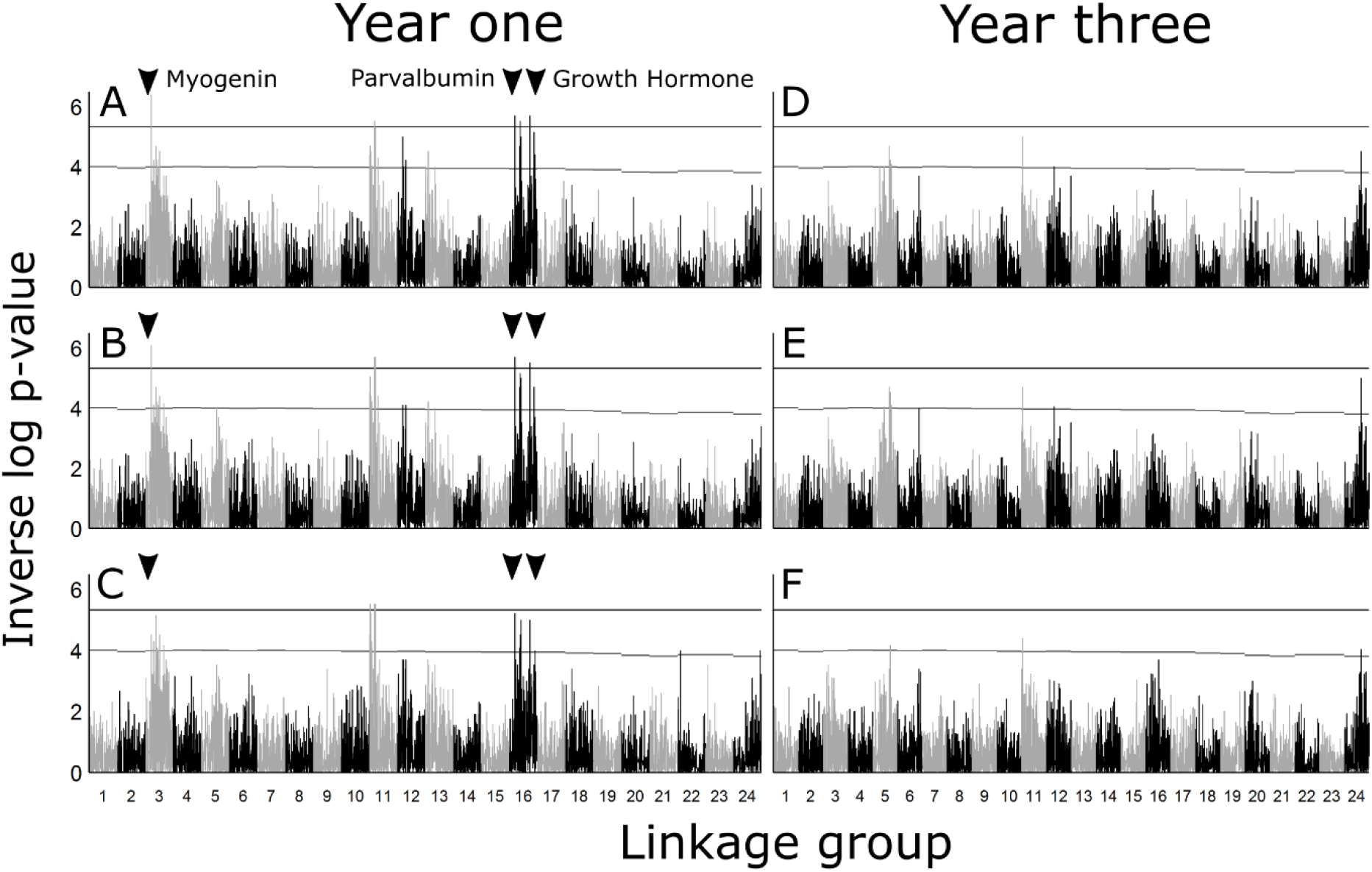
QTL scans at year one and year three for fork length (A+D), peduncle length (B+E), and weight (C+F) in the F_2_ generation of *Chrysophrys auratus.* The upper and lower line on each graph indicate the 95% genome-wide and linkage group-wide significance levels, respectively. Samples sizes varied for each subplot dependent on available data: A = 539, B = 539, C = 532, D = 312, E = 312, F = 323. The positions of three candidate genes known to be involved in growth are also indicated (a full list of candidate genes reviewed is shown in Supplementary Table 2).

### Candidate genes

The base-pair position on the genome scaffolds were found for all 13 candidate genes (Supplementary table 2) including growth hormone, growth hormone receptor, growth hormone receptor type 1, growth hormone receptor type 2, insulin like growth factor 1, insulin like growth factor 2, myogenic factor 1, myogenic factor 2, myogenic regulatory factor 4, myogenic regulatory factor 6, myogenin, myostatin, and parvalbumin. Based on the largest exon, all genes exhibited high base-pair similarity with the target genome position (88.7 to 99.3%). Of the candidate genes investigated, growth hormone, myogenin, and paravalbumin were located on linkage groups containing genome-wide significant QTLs (Figure 5; these were located within 13.2, 2.6, and 5.0 cM of a genome-wide significant QTL peak, respectively.

## DISCUSSION

We assembled the first chromosome level linkage map for the Australian snapper. Proof checking against our de novo genome assembly indicated that the linkage groups were of high quality. QTL mapping revealed eight markers on three linkage groups that were significantly associated with growth rate. Three candidate genes for growth rate were located on the same linkage groups as these QTL. These genomic resources will be used to advance the selective breeding programme currently underway on the study population and will form the basis of further genomic investigation in snapper.

Linkage maps have been used extensively in conjunction with QTL mapping studies to arrange markers into the order they appear in the target species genome (Boulton et al., 2011; Cnaani et al., 2004; Greenwood et al., 2011). The usefulness of a linkage map is determined by marker density and overall accuracy and precision. Historically, most first generation linkage maps in fish have been constructed with just a few hundred markers and did not have genome sequences available to evaluate their accuracy (Castaño-Sánchez et al., 2010). However, a few recent examples have increased the number of markers (Castaño-Sánchez et al., 2010; Ninwichian et al., 2012) and some have also begun to utilize available genome data for checking the linkage map accuracy and/or even check genome assemblies (Tsai et al., 2016; Wang et al., 2017). The snapper linkage map had ~11K markers for a 738 Mb genome (Figure 2), compared with ~96K markers in Atlantic salmon (2.97 Gb genome) – perhaps the most comprehensive fish linkage map to date (Tsai et al., 2016). The average correlation between the largest genome scaffolds (bp) and linkage map (cM) was 0.74. Comparatively, the results for the recently constructed salmon linkage map were 0.81 for the male map and 0.92 for the female map. The correlation between the linkage map and genome scaffolds is illustrated for the four largest scaffolds (Figure 3). While the correlation is clear, it is also apparent that at a fine scale (< 5 cM intervals) there is some variation around the exact placement of SNPs. In many cases this variation is probably the result of inherent precision limitations in the dataset (sample size + number of recombination events), but some of this variation could also be the result of differential recombination patterns across the genome (as observed in Roesti et al., 2013).

Using the newly constructed linkage map and available genome scaffolds, we were able to calculate the sex-specific recombination rates for snapper (female = 3.28 cM/Mb, male = 1.93 cM/Mb). This reflects similar observations in other fish species, with females often having higher recombination rates than males (Castaño-Sánchez et al., 2010; Kucuktas et al., 2009; Tsai et al., 2016). Overall, the recombination rates in this study are similar for those found in several fish species including stickleback *(Gasterosteus aculeatus,* 3.11 cM/Mb, (Roesti et al., 2013)), Asian seabass *(Lates calcarifer,* 2.4-2.8 cM/Mb, (Wang et al., 2017)), and channel catfish *(Ictalurus punctatus,* 2.6 cM/Mb, (Li et al., 2015)). Recent work has indicated that the different recombination rates for male and female fish may be related to the processes that occur during the formation of the egg or sperm (Theodosiou et al., 2016).

While dense, accurate, and precise linkage maps are useful for QTL mapping, very large pedigrees are needed to construct them; studies can require many thousands of individuals and families to get precise marker placement (He et al., 2011). Reference genomes provide an alternative way to position markers at their exact base pair location, but accuracy does depend on the stage and quality of the genome assembly (Berner et al., 2014). However, when a species does not have a fully assembled reference genome, linkage maps can prove valuable tools for determining the position of markers and aiding the arrangement of large genome scaffolds into new reference genomes (Fierst, 2015). Future work in snapper would benefit greatly from the combined use of the linkage map and available genome scaffolds to position SNPs more precisely. To do this, the linkage map could be used to position large genome scaffolds into chromosomes. From here, the arrangement of markers on the scaffold could be used to arrange the markers in the linkage map at a finer scales.

The target trait for this study was growth rate (Figure 1), which was measured using peduncle length, fork length, and weight (Table 1). Growth rate is one of the primary targets for selective breeding programmes because it relates directly to production output. In fish, it typically has a moderate degree of heritability, and in the current population was shown to be approximately ~0.26 and ~0.11 in year one and year three, respectively (Ashton et al., 2018). Other factors that can affect growth rate include feed amounts, fish density in tanks, and tank design (size, aeration, water flow). We attempted to control for these factors in the current study by standardizing the conditions between tanks and by standardizing measures from each tank before using them in QTL analysis.

Genome-wide significant QTLs were found in the first year, but not in the third year, which could indicate that the genetic basis of growth in early life is lost as the fish age (Figure 5, Table 2, Supplementary table 1). This is also probably in part a result of the decreased sample size from year one to year three, but may also reflect the lower heritability of growth rate in year three for the sample population. The lower sample size in year three was the result of natural mortality over the course of the study. Quantitative trait loci, of genome-wide significance, were highly shared among the three measures (peduncle length, fork length, and weight), which reflects that these are all measures of the same underlying trait (growth rate). The effect size ranged from 0.04 to 0.1 for individual QTLs, which is similar to growth QTLs observed in other species including Atlantic salmon *(Salmo salar.* 0.06 to 0.08, (Besnier et al., 2015)), tilapia *(Oreochromis niloticus:* 0.06 to 0.19, (Cnaani et al., 2004)), chinook salmon *(Oncorhynchus tshawytscha:* 0.14 to 0.33, (Everett and Seeb, 2014)), brill *(Scophthalmus rhombus:* 0.08 to 0.12, (Houston et al., 2009)), and catfish *(Ictalurus furcatus:* 0.01 to 0.23, (Hutson et al., 2014)). The identification of multiple QTLs affecting growth rate indicates a polygenic basis for growth rate. The effect size in snapper is on the lower side compared with that observed in these other studies, which could suggest that some other non-genetic factor is playing a larger role in determining the growth rate of this species in this specific culture environment.

Determining precision of QTL placement is an important step in the QTL mapping process as it provides useful information about where variants responsible for an observed QTL signal (e.g. candidate genes or causative alleles) are likely to be located. If high rates of linkage are present between markers, a confidence interval for the QTL region can be estimated - as seen in R/QTL (Broman et al., 2003). However, in the current study, pairwise correlation between markers (linkage) across the linkage map indicated very low linkage between markers over even relatively short distances (< 5 cM) (Figure 4). This is most obvious when comparing the linkage observed within the single largest family to that observed in the entire dataset (Figure 4). However, it is worth noting that on linkage groups with genome-wide significant QTLs, there does appear to be a number of markers surrounding each QTL that are responding to the QTL signal. As such, it seems likely that there is some linkage between markers at a fine scale (< 5 cM), but that this may be obscured by the low precision of marker placement on the linkage map. If true, the optimal way to get more precise placement of QTL regions will be improved SNP positioning using either a second improved iteration of the linkage map, the genome assembly, or a combination of both these resources. Until this is done it is likely that causative genetic variations underlying the QTL signals will be within this 5 cM scale.

Previous studies have outlined a range of genes and molecular networks that are thought to be candidates for further investigation in teleost species (De-Santis and Jerry, 2007). While the QTL regions in the current study were too large to identify a definitive link between the QTL signals and candidate genes, we did locate the position of candidate growth genes close to putative QTLs (Supplementary table 2). This was possible using the available genome sequence data to link gene positions back to their nearest markers. Central to growth rate in most species is the somatotropic axis, which consists of the growth hormone releasing hormone (GHRH), growth hormone inhibiting hormone (GHIH), growth hormone (GH), and insulin-like growth factors (IGF-1 and –II) (De-Santis and Jerry, 2007). Of these, growth hormone and insulin-like growth factor I and II were able to be mapped to the linkage map in the current study. Growth hormone was located near (within 13.2 cM) a QTL of genome-wide significance. In *Sparus aurata,* a close relative of *C. auratus,* a microsatellite repeat in the promoter region has previously been implicated for differences in growth rate (Almuly et al., 2000). This gene would be a good candidate in *C. auratus* because it is close to a QTL in the current study, and the causative microsatellite has been observed in a range of teleost species. Myogenic regulatory factors (myogenin, MyoD, myf-5, and myf-6) are another set of potential candidate genes (De-Santis and Jerry, 2007). These regulatory factors have been implicated in growth in terrestrial vertebrates, but not in fish species. In this study, the myogenin gene was located on linkage group 3 within 2.6 cM of a genome-wide significant QTL. In pigs, a polymorphism in the promoter region of myogenin was found to account for up to 5.8% of differences in weight (te Pas et al., 1999), but no research has investigated its effect in teleost species. A final candidate gene that was positioned close (within 5cM) to a putative growth rate QTL was parvalbumin. A mutation in the promoter region of this gene was found to be involved in weight differences in the finfish species *Lates calcarifer.*

### Future directions

While this study can confidently place the QTL in ~5 cM regions, further work is needed to define the QTL regions more precisely. Improving the precision of these regions would aid fine-mapping and further characterization of the described QTLs and related candidate genes. Improved precision should be possible in the near future using the assembled genome that is currently being developed at Plant & Food Research. Future work should also aim to detect possible sex-linked markers, to identify regions associated with sex determination, and to investigate sex-specific recombination patterns across the genome. While sex-specific information was not investigated in the current study, this is an area of particular interest in snapper and the data from this study could be used to further investigate it. In conclusion, this study provides valuable genetic and genomic resources for future evolutionary studies and aquaculture breeding programmes in this and related species.

## AUTHOR CONTRIBUTIONS

Conceptualization, DTA, PAR, and MW; Methodology, DTA; Investigation, DTA; Data Curation, DTA; Writing – Original Draft, DTA; Writing – Review & Editing, DTA, PAR, and MW; Funding Acquisition, MW; Supervision, PAR and MW.

## ACKNOWLEDGEMENTS

We would like to acknowledge all the PFR staff who have been involved with this work in particular Peter Jaksons, Ross Crowhurst, and Elena Hilario.

## SUPPLEMENTARY

**Supplementary figure 1.**
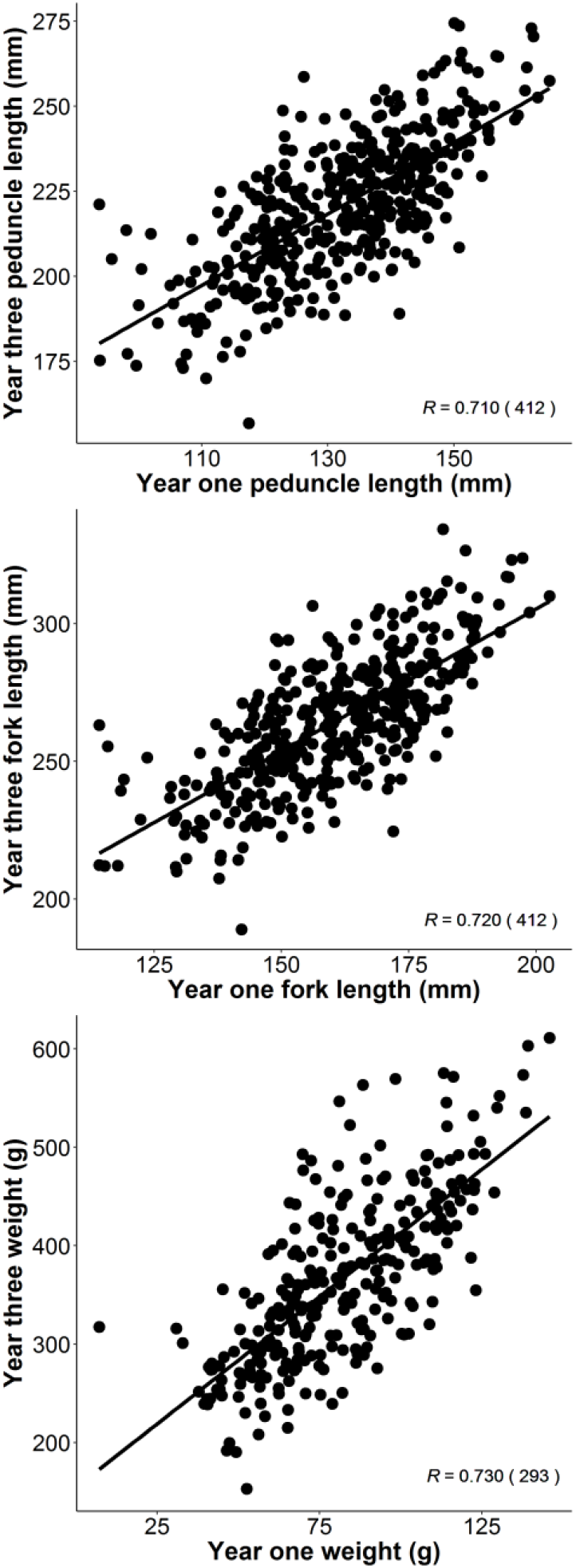
Correlation between fork length, peduncle length, and weight between individuals in year one and the same individuals in year three.

**Supplementary Table 1.**
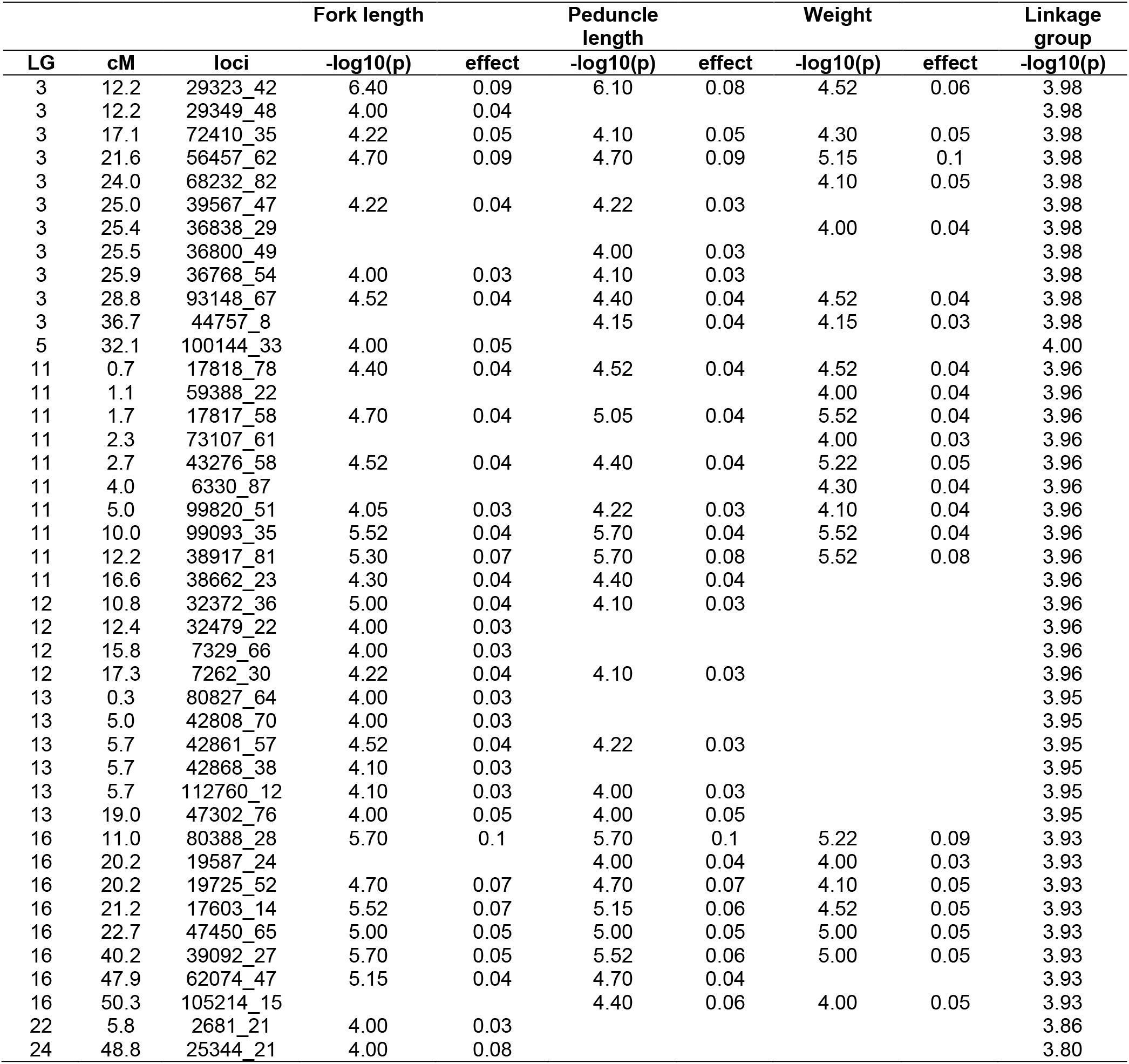
A complete list of chromosome and genome wide significant QTLs from the current study including linkage group (LG), centimorgan position (cM), marker name (loci), significance, and effect size. The genome-wide significance limit -log10(p) was 5.33.

**Supplementary table 2.** Candidate gene information and position on the linkage map. An asterix * indicates that the gene was located on a linkage group also containing a genome-wide significant QTL for growth rate.

| Gene | Species | Accession # | Type | LG | cM | bp | % match |
| --- | --- | --- | --- | --- | --- | --- | --- |
| growth-hormone* | *Pagrus major* | AB904715.1 | DNA | 16 | 53.2 | 147 | 99.3 |
| myostatin | *Pagrus major* | AY965686.1 | DNA | 6 | 38.3 | 403 | 99.0 |
| growth-hormone-receptor | *Epinephelus coioides* | KR269817.1 | DNA | 9 | 58.3 | 769 | 88.7 |
| growth-hormone-receptor-type-I | *Sparus aurata* | AH014067.4 | DNA | 9 | 58.3 | 760 | 95.4 |
| growth-hormone-receptor-type-II | *Sparus aurata* | AH014068.4 | DNA | 18 | 50.3 | 875 | 94.1 |
| myogenin* | *Sparus aurata* | EF462192.1 | DNA | 3 | 9.6 | 534 | 96.8 |
| myogenic-factor-MYOD1 | *Sparus aurata* | AF478568.1 | DNA | 7 | 12.6 | 591 | 97.0 |
| myogenic-factor-MYOD2 | *Sparus aurata* | AF478569.1 | DNA | 14 | 39 | 546 | 96.5 |
| myogenic-regulatory-factor-4 | *Epinephelus coioides* | KR269828.1 | DNA | 15 | 25.2 | 510 | 93.8 |
| myogenic-regulatory-factor-6 | *Sparus aurata* | JN034421.1 | mRNA | 15 | 25.2 | 521 | 95.6 |
| insulin-like-growth-factor-I | *Sparus aurata* | DQ118098.1 | DNA | 15 | 25.2 | 186 | 99.5 |
| insulin-like-growth-factor-II | *Epinephelus coioides* | KR269813.1 | DNA | 7 | 23.4 | 240 | 96.7 |
| parvalbumin* | *Sparus aurata* | GU060310.1 | mRNA | 16 | 15.2 | 311 | 97.7 |

## LITERATURE CITED

Abecasis, G.R., Cardon, L.R., Cookson, W.O., 2000. A general test of association for quantitative traits in nuclear families. American Journal of Human Genetics 66, 279–292. https://doi.org/10.1086/302698

Almuly, R., Cavari, B., Ferstman, H., Kolodny, O., Funkenstein, B., 2000. Genomic structure and sequence of the gilthead seabream (*Sparus aurata*) growth hormone-encoding gene: identification of minisatellite polymorphism in intron I. Genome 43, 836–845.

Ashton, D.T., Hilario, E., Jaksons, P., Wellenreuther, M., 2018. Genetic diversity and heritability of economically important traits in captive Australasian snapper (*Chrysophrys auratus*). in prep.

Barria, A., Christensen, K.A., Correa, K., Jedlicki, A., Lhorente, J.P., Davidson, W., Yáñez, J.M., 2017. Genome-Wide Association Study And Genomic Predictions For Resistance Against Piscirickettsia salmonis In Coho Salmon (*Oncorhynchus kisutch*) Using ddRAD Sequencing. bioRxiv 124099. https://doi.org/10.1101/124099

Berner, D., Moser, D., Roesti, M., Buescher, H., Salzburger, W., 2014. Genetic Architecture of Skeletal Evolution in European Lake and Stream Stickleback. Evolution 68, 1792–1805. https://doi.org/10.1111/evo.12390

Besnier, F., Glover, K.A., Lien, S., Kent, M., Hansen, M.M., Shen, X., Skaala, Ø., 2015. Identification of quantitative genetic components of fitness variation in farmed, hybrid and native salmon in the wild. Heredity 115, 47.

Bettembourg, M., Dardou, A., Audebert, A., Thomas, E., Frouin, J., Guiderdoni, E., Ahmadi, N., Perin, C., Dievart, A., Courtois, B., 2017. Genome-wide association mapping for root cone angle in rice. Rice 10, 45. https://doi.org/10.1186/s12284-017-0184-z

Bhatta, S., Iwai, T., Miura, C., Higuchi, M., Shimizu-Yamaguchi, S., Fukada, H., Miura, T., 2012. Gonads directly regulate growth in teleosts. Proceedings of the National Academy of Sciences 109, 11408–11412. https://doi.org/10.1073/pnas.1118704109

Boulton, K., Massault, C., Houston, R.D., de Koning, D.J., Haley, C.S., Bovenhuis, H., Batargias, C., Canario, A.V.M., Kotoulas, G., Tsigenopoulos, C.S., 2011. QTL affecting morphometric traits and stress response in the gilthead seabream (*Sparus aurata*). Aquaculture 319, 58–66. https://doi.org/10.1016/j.aquaculture.2011.06.044

Braasch, I., Peterson, S.M., Desvignes, T., McCluskey, B.M., Batzel, P., Postlethwait, J.H., 2015. A new model army: Emerging fish models to study the genomics of vertebrate Evo-Devo. Journal of Experimental Zoology Part B: Molecular and Developmental Evolution 324, 316–341. https://doi.org/10.1002/jez.b.22589

Broman, K.W., Wu, H., Sen, S., Churchill, G.A., 2003. R/qtl: QTL mapping in experimental crosses. Bioinformatics 19, 889–890.

Castaño-Sánchez, C., Fuji, K., Ozaki, A., Hasegawa, O., Sakamoto, T., Morishima, K., Nakayama, I., Fujiwara, A., Masaoka, T., Okamoto, H., Hayashida, K., Tagami, M., Kawai, J., Hayashizaki, Y., Okamoto, N., 2010. A second generation genetic linkage map of Japanese flounder (*Paralichthys olivaceus*). BMC Genomics 11, 554. https://doi.org/10.1186/1471-2164-11-554

Catchen, J., Hohenlohe, P.A., Bassham, S., Amores, A., Cresko, W.A., 2013. Stacks: an analysis tool set for population genomics. Molecular Ecology 22, 3124–3140. https://doi.org/10.1111/mec.12354

Chen, H., Semagn, K., Iqbal, M., Moakhar, N.P., Haile, T., N’Diaye, A., Yang, R.-C., Hucl, P., Pozniak, C., Spaner, D., 2017. Genome-wide association mapping of genomic regions associated with phenotypic traits in Canadian western spring wheat. Molecular Breeding 37, 141. https://doi.org/10.1007/s11032-017-0741-6

Cnaani, A., Zilberman, N., Tinman, S., Hulata, G., Ron, M., 2004. Genome-scan analysis for quantitative trait loci in an F_2_ tilapia hybrid. Molecular Genetics and Genomics 272, 162–172. https://doi.org/10.1007/s00438-004-1045-1

Cros, D., Bocs, S., Riou, V., Ortega-Abboud, E., Tisné, S., Argout, X., Pomiès, V., Nodichao, L., Lubis, Z., Cochard, B., Durand-Gasselin, T., 2017. Genomic preselection with genotyping-by-sequencing increases performance of commercial oil palm hybrid crosses. BMC Genomics 18, 839. https://doi.org/10.1186/s12864-017-4179-3

Dekkers, J.C.M., 2012. Application of Genomics Tools to Animal Breeding. Current Genomics 13, 207–212. https://doi.org/10.2174/138920212800543057

De-Santis, C., Jerry, D.R., 2007. Candidate growth genes in finfish — Where should we be looking? Aquaculture 272, 22–38. https://doi.org/10.1016/j.aquaculture.2007.08.036

Everett, M.V., Seeb, J.E., 2014. Detection and mapping of QTL for temperature tolerance and body size in Chinook salmon (*Oncorhynchus tshawytscha*) using genotyping by sequencing. Evolutionary Applications 7, 480–492. https://doi.org/10.1111/eva.12147

Fierst, J.L., 2015. Using linkage maps to correct and scaffold de novo genome assemblies: methods, challenges, and computational tools. Frontiers in Genetics 6. https://doi.org/10.3389/fgene.2015.00220

Greenwood, A.K., Jones, F.C., Chan, Y.F., Brady, S.D., Absher, D.M., Grimwood, J., Schmutz, J., Myers, R.M., Kingsley, D.M., Peichel, C.L., 2011. The genetic basis of divergent pigment patterns in juvenile threespine sticklebacks. Heredity 107, 155. https://doi.org/10.1038/hdy.2011.1

Handeland, S.O., Imsland, A.K., Stefansson, S.O., 2008. The effect of temperature and fish size on growth, feed intake, food conversion efficiency and stomach evacuation rate of Atlantic salmon post-smolts. Aquaculture 283, 36–42. https://doi.org/10.1016/j.aquaculture.2008.06.042

He, C., Weeks, D.E., Buyske, S., Abecasis, G.R., Stewart, W.C., Matise, T.C., 2011. Enhanced genetic maps from family-based disease studies: population-specific comparisons. BMC Medical Genetics 12, 15. https://doi.org/10.1186/1471-2350-12-15

Hilario, E., 2015. The Restriction Enzyme Target Approach to Genotyping by Sequencing (GBS), in: Plant Genotyping, Methods in Molecular Biology. Humana Press, New York, NY, pp. 271–279. https://doi.org/10.1007/978-1-4939-1966-6_20

Houston, R.D., Bishop, S.C., Hamilton, A., Guy, D.R., Tinch, A.E., Taggart, J.B., Derayat, A., McAndrew, B.J., Haley, C.S., 2009. Detection of QTL affecting harvest traits in a commercial Atlantic salmon population. Animal Genetics 40, 753–755. https://doi.org/10.1111/j.1365-2052.2009.01883.x

Hutson, A.M., Liu, Z., Kucuktas, H., Umali-Maceina, G., Su, B., Dunham, R.A., 2014. Quantitative trait loci map for growth and morphometric traits using a channel catfish × blue catfish interspecific hybrid system. Journal of Animal Science 92, 1850–1865. https://doi.org/10.2527/jas.2013-7191

Kalinowski, S.T., Taper, M.L., Marshall, T.C., 2007. Revising how the computer program cervus accommodates genotyping error increases success in paternity assignment. Molecular Ecology 16, 1099–1106. https://doi.org/10.1111/j.1365-294X.2007.03089.x

Kearse, M., Moir, R., Wilson, A., Stones-Havas, S., Cheung, M., Sturrock, S., Buxton, S., Cooper, A., Markowitz, S., Duran, C., Thierer, T., Ashton, B., Meintjes, P., Drummond, A., 2012. Geneious Basic: An integrated and extendable desktop software platform for the organization and analysis of sequence data. Bioinformatics 28, 1647–1649. https://doi.org/10.1093/bioinformatics/bts199

Kucuktas, H., Wang, S., Li, P., He, C., Xu, P., Sha, Z., Liu, H., Jiang, Y., Baoprasertkul, P., Somridhivej, B., Wang, Y., Abernathy, J., Guo, X., Liu, L., Muir, W., Liu, Z., 2009. Construction of Genetic Linkage Maps and Comparative Genome Analysis of Catfish Using Gene-Associated Markers. Genetics 181, 1649–1660. https://doi.org/10.1534/genetics.108.098855

Lehner, B., 2013. Genotype to phenotype: lessons from model organisms for human genetics. Nature Reviews Genetics 14, 168. https://doi.org/10.1038/nrg3404

Li, Y., Liu, S., Qin, Z., Waldbieser, G., Wang, R., Sun, L., Bao, L., Danzmann, R.G., Dunham, R., Liu, Z., 2015. Construction of a high-density, high-resolution genetic map and its integration with BAC-based physical map in channel catfish. DNA Research 22, 39–52. https://doi.org/10.1093/dnares/dsu038

Lynch, M., Walsh, B., 1998. Genetics and analysis of quantitative traits. Sinauer Sunderland, MA.

Miller, S., Harley, J., 2006. Zoology. McGraw-Hill Higher Education.

Ninwichian, P., Peatman, E., Liu, H., Kucuktas, H., Somridhivej, B., Liu, S., Li, P., Jiang, Y., Sha, Z., Kaltenboeck, L., Abernathy, J.W., Wang, W., Chen, F., Lee, Y., Wong, L., Wang, S., Lu, J., Liu, Z., 2012. Second-Generation Genetic Linkage Map of Catfish and Its Integration with the BAC-Based Physical Map. G3: Genes|Genomes|Genetics 2, 1233–1241. https://doi.org/10.1534/g3.112.003962

Park, I.-S., Gil, H.W., Lee, T.H., Nam, Y.K., Kim, D.S., 2016. Comparative Study of Growth and Gonad Maturation in Diploid and Triploid Marine Medaka, *Oryzias dancena*. Development & Reproduction 20, 305–314. https://doi.org/10.12717/DR.2016.20.4.305

Pértille, F., Moreira, G.C.M., Zanella, R., Nunes, J. de R. da S., Boschiero, C., Rovadoscki, G.A., Mourão, G.B., Ledur, M.C., Coutinho, L.L., 2017. Genome-wide association study for performance traits in chickens using genotype by sequencing approach. Scientific Reports 7. https://doi.org/10.1038/srep41748

Purcell, S., Neale, B., Todd-Brown, K., Thomas, L., Ferreira, M.A.R., Bender, D., Maller, J., Sklar, P., de Bakker, P.I.W., Daly, M.J., Sham, P.C., 2007. PLINK: A Tool Set for Whole-Genome Association and Population-Based Linkage Analyses. American Journal of Human Genetics 81, 559–575.

R Core Team, 2013. R: A language and environment for statistical computing. R Foundation for Statistical Computing, Vienna, Austria.

Rastas, P., Calboli, F.C.F., Guo, B., Shikano, T., Merilä, J., 2016. Construction of Ultradense Linkage Maps with Lep-MAP2: Stickleback F_2_ Recombinant Crosses as an Example. Genome Biology and Evolution 8, 78–93. https://doi.org/10.1093/gbe/evv250

Roesti, M., Moser, D., Berner, D., 2013. Recombination in the threespine stickleback genome—patterns and consequences. Molecular Ecology 22, 3014–3027. https://doi.org/10.1111/mec.12322

Savolainen, O., Lascoux, M., Merilä, J., 2013. Ecological genomics of local adaptation. Nature Reviews Genetics 14, 807. https://doi.org/10.1038/nrg3522

Spielman, R.S., Ewens, W.J., 1996. The TDT and other family-based tests for linkage disequilibrium and association. American Journal of Human Genetics 59, 983–989.

te Pas, M.F., Soumillion, A., Harders, F.L., Verburg, F.J., van den Bosch, T.J., Galesloot, P., Meuwissen, T.H., 1999. Influences of myogenin genotypes on birth weight, growth rate, carcass weight, backfat thickness, and lean weight of pigs. Journal of Animal Science 77, 2352–2356.

Theodosiou, L., McMillan, W.O., Puebla, O., 2016. Recombination in the eggs and sperm in a simultaneously hermaphroditic vertebrate. Proceedings of the Royal Society B: Biological Sciences 283. https://doi.org/10.1098/rspb.2016.1821

Tsai, H.Y., Hamilton, A., Guy, D.R., Tinch, A.E., Bishop, S.C., Houston, R.D., 2015. The genetic architecture of growth and fillet traits in farmed Atlantic salmon (*Salmo salar*). BMC Genetics 16, 51. https://doi.org/10.1186/s12863-015-0215-y

Tsai, H.Y., Robledo, D., Lowe, N.R., Bekaert, M., Taggart, J.B., Bron, J.E., Houston, R.D., 2016. Construction and Annotation of a High Density SNP Linkage Map of the Atlantic Salmon (*Salmo salar*) Genome. G3: Genes|Genomes|Genetics 6, 2173–2179. https://doi.org/10.1534/g3.116.029009

Wang, C., 2009. Quantitative genetic estimates of growth-related traits in the common carp (*Cyprinus carpio L.*): A review. Frontiers of Biology in China 4, 298–304. https://doi.org/10.1007/s11515-009-0031-8

Wang, L., Bai, B., Liu, P., Huang, S.Q., Wan, Z.Y., Chua, E., Ye, B., Yue, G.H., 2017. Construction of high-resolution recombination maps in Asian seabass. BMC Genomics 18. https://doi.org/10.1186/s12864-016-3462-z

Ye, B., Wan, Z., Wang, L., Pang, H., Wen, Y., Liu, H., Liang, B., Lim, H.S., Jiang, J., Yue, G., 2017. Heritability of growth traits in the Asian seabass (*Lates calcarifer*). Aquaculture and Fisheries 2, 112–118. https://doi.org/10.1016/j.aaf.2017.06.001

